# Expressed *vomeronasal type-1 receptors* (*V1rs*) in bats uncover conserved mechanisms of social chemical signaling

**DOI:** 10.1101/293472

**Authors:** Laurel R. Yohe, Kalina T. J. Davies, Stephen J. Rossiter, Liliana M. Dávalos

**Author notes:** **Corresponding Author:** Laurel R. Yohe 650 Life Sciences, Stony Brook, NY 11794; LRY.

## Abstract

In mammals, social and reproductive behaviors are mediated by chemical cues encoded by hyperdiverse families of receptors expressed in the vomeronasal organ. Between species, the number of intact receptors can vary by orders of magnitude. However, the evolutionary processes behind variation in receptor number, and also its link to fitness-related behaviors are not well understood. From vomeronasal transcriptomes, we discovered the first evidence of intact *vomeronasal type-1 receptor* (*V1r*) genes in bats, and we tested whether putatively functional bat receptors were orthologous to those of related taxa, or whether bats have evolved novel receptors. We found that *V1r*s in bats and show high levels of orthology to those of their relatives, as opposed to lineage-specific duplications, and receptors are under purifying selection. Despite widespread vomeronasal organ loss in bats, *V1r* copies have been retained for >65 million years. The highly conserved nature of bat *V1r*s challenges our current understanding of mammalian *V1r* function and suggest roles other than conspecific recognition or mating initiation in social behavior.

---

Nearly all mammals can perceive pheromones—broadly construed as any olfactory cue excreted from individuals of a different species or conspecific (Silva & Antunes 2017)—though there is great variation in the genetic detection mechanism and morphological structures involved (Young et al. 2010; Meisami & Bhatnagar 1998; Grus et al. 2007). Mammalian pheromone detection, or vomerolfaction (Cooper & Burghardt 1992), mediates key social and reproductive behaviors including mating and courtship, parental care, conspecific identification, and territoriality (Liberles 2014). Pheromone detection occurs in the vomeronasal organ, composed of a cluster of sensory neurons in the nasal anterior that express ultrasensitive G-protein coupled receptors (e.g. vomeronasal type-1 receptors [V1Rs], vomeronasal type-2 receptors [V2Rs]). These receptors bind to the pheromones (Ibarra-Soria et al. 2014), and trigger a signaling cascade that activates the Transient receptor potential cation channel 2 (Trpc2) ion channel resulting in depolarization, so the cue can be processed by the brain (Mast et al. 2010). However, pinpointing which of the hundreds of receptors mediates a given behavior is challenging. A comparative approach can narrow down the scope of functional characterization, as understanding the gene history and molecular evolution across divergent lineages can help determine which receptors are relevant to particular species. Here we analyze the diversity of mammalian *V1r*s —focusing particularly on bats—and infer the processes responsible for their evolutionary history. We concentrate primarily on *V1r*s, as they show the greatest variation in number of genes across species of any mammalian gene family (Grus et al. 2007; Young et al. 2010), and dominate among vomeronasal receptors in placental mammals (Silva & Antunes 2017).

Despite its importance in fitness-related behaviors, the vomeronasal organ is vestigial in a few clades including several aquatic mammals, catarrhine primates, and many bats (Bhatnagar & Meisami 1998; Zhang & Webb 2003; Yohe et al. 2017; Yu et al. 2010). The relaxation of selection in these lineages has led to pseudogenization of many elements of the molecular pathways involved in pheromone detection and transduction (Yohe et al. 2017; Yu et al. 2010; Zhang & Webb 2003; Young et al. 2010; Zhao et al. 2011), and losses may be related to shifts to underwater or diurnal niches. No explanation has emerged, however, for variation in the maintenance of the vomeronasal system of bats, as more than a dozen independent functional losses in *Trpc2* gene function seem unrelated to either the evolution of flight, or of other specialized senses (Yohe et al. 2017).

*V1r*s play a role in species-specific behaviors (Ibarra-Soria et al. 2014; Grus & Zhang 2004), and may even play a role in speciation. For example, in rodents, orthologous receptors vary among species and subspecies, with less than 20% of genes shared between mouse and rat (Zhang et al. 2007; Park et al. 2011; Wynn et al. 2012). Duplications of *V1r*s prior to the diversification of lemurs and lorises expanded the number of intact *V1r*s by an order of magnitude (Yoder et al. 2014; Yoder & Larsen 2014), perhaps promoting strepsirrhine diversification as they colonized Madagascar. Like other chemosensory genes, *V1r*s evolve via a birth-death process by which gene copies frequently duplicate and pseudogenize over time (Nei & Rooney 2005). This birth-death process genearates great variance in receptor numbers across species; for example, there are well over 200 *V1r*s in the platypus and mouse lemurs, fewer than 10 intact *V1r*s in catarrhine primates, and none were detected in either the bottlenose dolphin or the two species of bats previously analyzed (Young et al. 2010). Attempts to explain this variance have linked *V1r* numbers to nocturnality (Wang et al. 2010), but correlating numbers of receptors to functional ecology fails to address the evolutionary history of *V1r* repertoires. Here we trace the phylogenetic history of each bat *V1r* gene and infer its orthology to determine whether each *V1r* is shared among divergent mammals, or instead unique to a species or clade. Because *V1r*s have been shown to mediate species-specific behaviors that may be related to species boundaries, we hypothesized that bat *V1r*s have evolved through lineage-specific duplications and perhaps served as a key innovation that facilitated speciation of the New World leaf-nosed bats (Phyllostomidae)—a species rich clade with diverse dietary adaptations and conserved a functional *Trpc2* (Yohe et al. 2017). Alternatively, *V1r*s may be conserved orthologs of non-bat lineages. As orthologous chemosensory genes of divergent species will have a higher probability of detecting a similar compound than will paralogs within a species (Adipietro et al. 2012), shared orthology among bats and non-bats could indicate that the receptor binds to similar ligands or mediate similar behaviors.

To test our hypotheses, we generated new transcriptomes from the vomeronasal organs of six species of phyllostomids (Table S1), and we combined these with data from published genomes of 13 additional species. Our data revealed at least one intact *V1r* in each transcriptome, thus providing the first evidence of transcribed *V1r*s in bats. The vampire bat (*Desmodus rotundus*) had eight distinct expressed *V1r*s, the most of any of the bat species we examined. We validated these receptor transcripts with the *V1r* sequences identified from the recently published vampire bat genome (Lisandra Zepeda Mendoza et al. 2017). With one exception, all transcribed *V1r*s were found among the 14 intact *V1r* sequences identified in the genome (Fig. S1).

We also characterized intact and pseudogenized *V1r*s from all other available bat genomes (14 in total), as well as the horse and the dog, two outgroup representatives within Laurasiatheria. In genome searches of bats from the suborder Yangochriptera, several intact *V1r*s were detected in *Miniopterus natalensis* and *Pteronotus parnellii*, two non-phyllostomid species previously shown to have an intact *Trpc2* gene (Fig. 1). However, we identified few intact *V1r*s in any other bat genome. An abundance of pseudogenized receptor genes were found in the exclusively Old World suborder Yinpterochiroptera, all of which have pseudogenized *Trpc2* genes (Fig. 1). Three species of yinpterochiropterans, of the 11 species predicted to lack a vomeronasal organ based on *Trpc2*, are an exception, with 1-2 *V1r*s with intact reading frames identified (Fig. 1). We also detected several fewer receptors (between 3-6 genes) from the horse and dog genomes than had been previously reported (Young et al. 2010). We emphasize, however, that the reported number of *V1r* genes per species should be considered a dynamic value and may change as genome assemblies and annotation methods improve.

**Figure 1.**
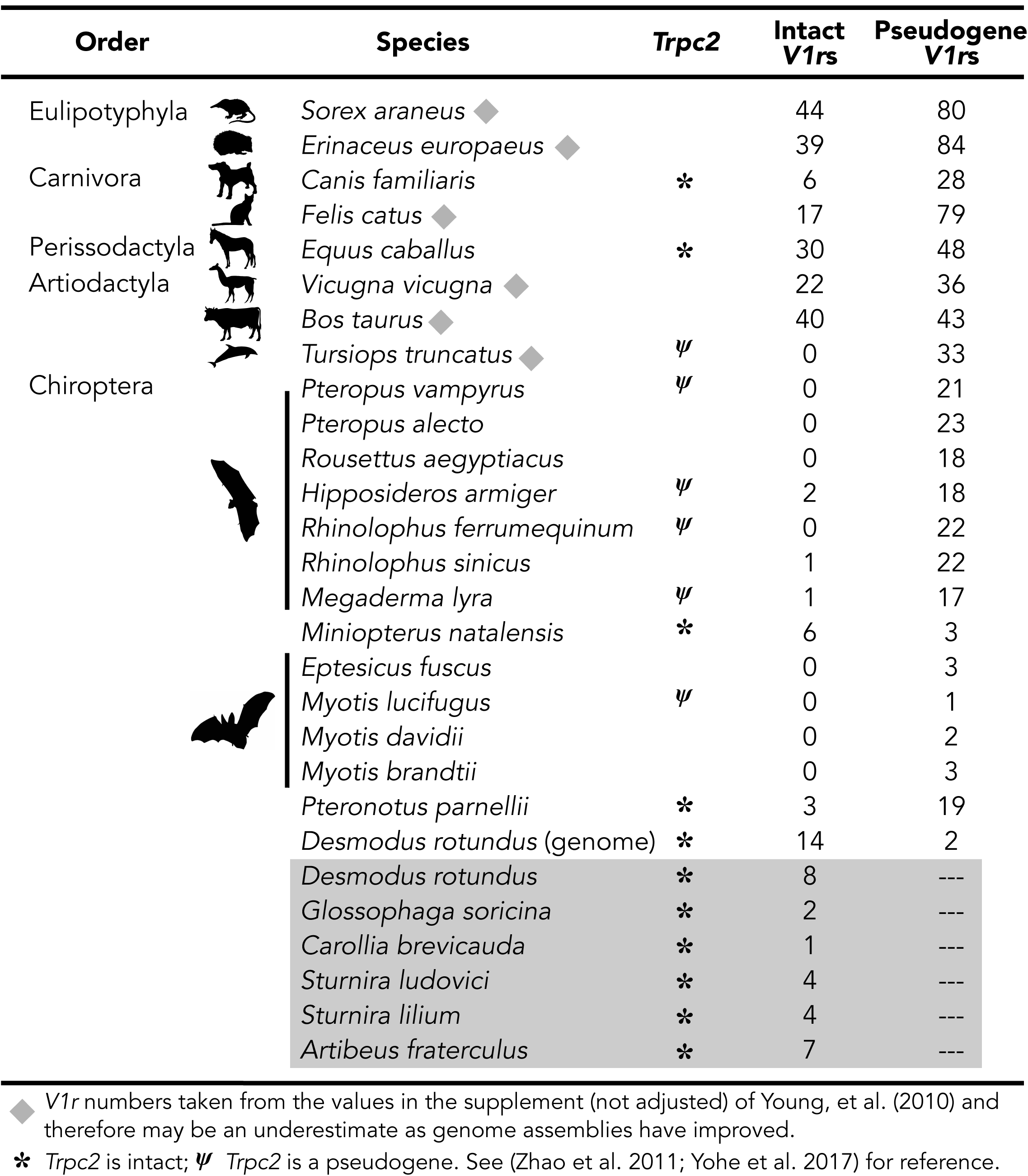
Number of intact and pseudogenized *V1r*s among laurasiatherians. *V1r*s from the transcriptome are highlighted in grey. The remaining species were characterized from available genomes. Pseudogenized *V1r*s are receptor genes with a frameshift or premature stop codon but with at least 650 base pairs. Vertical lines are bats that likely have a vestigial vomeronasal system, either based on morphology or *Trpc2*. Silhouettes are not to scale and were obtained from PhyloPic.

To determine orthologous gene groups (orthogroups) of *V1r*s, we reconstructed unrooted trees and identified genes forming monophyletic groups across different species (Ballesteros & Hormiga 2016). We pruned the gene tree into orthogroups while also allowing in-paralogs, genes within an orthogroup duplicated since a species diverged, to remain in the tree. Eighteen orthogroups were recovered, but five of these orthogroups contained only a single gene and many contained only two or three genes. Thus, we recovered a total of three orthogroups (Fig. 2A–C) informative for subsequent analyses of molecular evolution. An orthogroup of six bat-specific genes was recovered, but two distantly related bat and horse genes were excluded due to low bootstrap (Fig. 2D). There were no orthogroups with more than six genes that solely contained bats, suggesting all bats share orthologs with either the horse or dog lineage.

**Figure 2.**
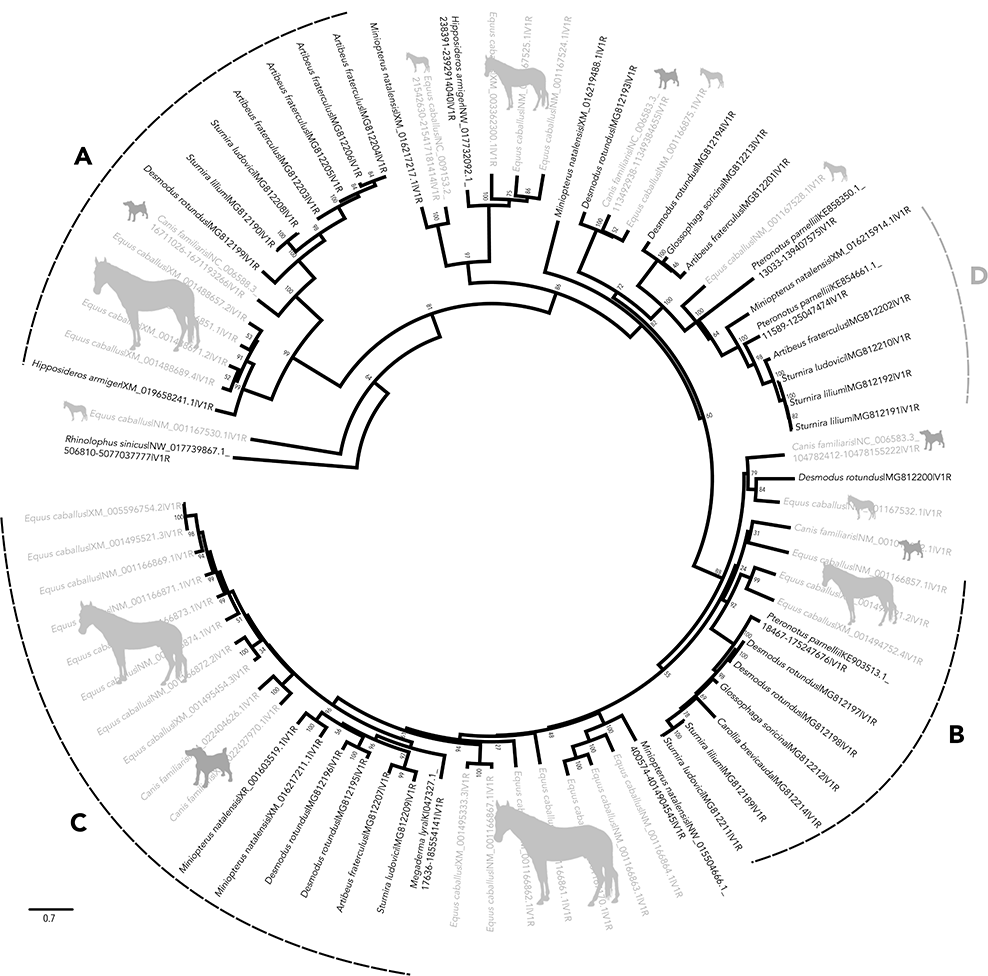
Codon model gene tree for intact *V1r*s identified from the vomeronasal organ transcriptomes of bats (black names), the few functional *V1r*s from bat genomes (also in black), and the genomes of *Equus caballus* and *Canis familiaris* (grey names). Node labels are bootstrap support values. Numbers on the tip label gene correspond the either the GenBank number (transcriptome data), RefSeq number, or genome location for newly identified genomic sequences in which no RefSeq number is available. Letter labels indicate orthogroups identified from the UPhO analysis that resulted in more than 6 taxa and included any non-bats. Orthogroup D is a bat-specific orthogroup that was not included in the selection analyses. Silhouettes were obtained from PhyloPic.

In mice, lemurs, and marsupials, considerable variation in *V1r* copies among species suggests vomerolfaction mediates species recognition, and possibly speciation (Yoder et al. 2014; Grus et al. 2005; Wynn et al. 2012). Although most bats with transcribed *V1r*s are found within the recently radiated New World leaf-nosed bats (Dumont et al. 2012), the small number of species-specific paralogs combined with the 100% orthology between bat receptors and those from the horse and dog (compared to ~10% orthology seen in mouse and 16% in rats (Ohara et al. 2009; Zhang et al. 2007; Grus & Zhang 2004)) together suggest that they play no role in species recognition. Hence, the low *V1r* diversity in bats implies an alternative function for these receptors. Comparisons in ruminants (cow, sheep, and goat) revealed conserved *V1r* repertoires with up to 70% orthology between species, but very little overlap with rodent *V1r* repertoires (Ohara et al. 2009). Like other laurasiatherians (Keller & Lévy 2012), bats display a high degree of orthology with their relatives. Such sequence conservation hints at function mediating innate behaviors common to all laurasiatherians, as the vomeronasal neurons that express *V1r*s are hard-wired to a common region of the brain responsible for similar instinctive behaviors (Bear et al. 2016), including mating, predator detection, and parental care. Although the receptors may differ in the compounds they bind as a result of amino acid differences among lineages; thus sequence conservation and orthology imply functions shared by all laurasiatherian species rather than species-specific roles.

To test for Darwinian selection in bat V1rs, we estimated the ratio of nonsynonymous to synonymous substitution rates (ω) for bats and compared to the background rate including genes from the horse and dog. First considering rates for the entire tree of intact *V1r*s, we found no significant difference between rates in bats and other species (Table 1 (PAML): χ^2^_(1)_ = 0.71 *P* = 0.40; Table 2 (RELAX): χ^2^_(1)_ = 0.05; *P* = 1.0), suggesting similar evolutionary processes are shaping the *V1r* repertoires of bats and non-bats. Nevertheless, rates of *V1r* molecular evolution are relatively high in both bats and their sampled relatives. Both across the entire phylogeny of intact receptors and within orthogroups (Table 1, 2), there were at least 48%, and sometimes as many as 62%, of codon sites evolving neutrally (ω = 1.0) in both bats and non-bats.

**Table 1.** Results from the PAML clade models. The grey box indicates the selected model or the null model not rejected based on the likelihood ratio test. Values for the site classes are ω estimates for each of the three site classes: purifying (ω_1_), neutral (ω_2_), and varying (ω_3_). The percentages in parentheses are the proportion of sites found within that respective site class.

| Model | lnL | np | K | TL | LR | p | $\omega$ site classes | | |
| --- | --- | --- | --- | --- | --- | --- | --- | --- | --- |
| | | | | | | | $\omega_1$ | $\omega_2$ | $\omega_3$ |
| Whole Tree |  |  |  |  |  |  |  |  |  |
| *M2a_rel (null)* | -36789 | 154 | 2.33 | 40.5 | --- | --- |  |  |  |
| $\omega_{\text{background}}$ | | | | | | | 0.15 (14%) | 1.0 (48%) | 0.48 (37%) |
| *Clade Model C* | -36789 | 155 | 2.33 | 40.5 | 0.71 | 0.40 |  |  |  |
| $\omega_{\text{background}}$ | | | | | | | 0.14 (14%) | 1.0 (49%) | 0.47 (37%) |
| $\omega_{\text{bats}}$ | | | | | | | 0.14 (14%) | 1.0 (49%) | 0.50 (37%) |
| Orthogroup A |  |  |  |  |  |  |  |  |  |
| *M2a_rel (null)* | -5593 | 28 | 2.67 | 4.43 | --- | --- |  |  |  |
| $\omega_{\text{background}}$ | | | | | | | 0.13 (25%) | 1.0 (62%) | 0.13 (12%) |
| *Clade Model C* | -5584 | 29 | 2.74 | 4.62 | 17.5 | 2.9e-5 |  |  |  |
| $\omega_{\text{background}}$ | | | | | | | 0.13 (37%) | 1.0 (62%) | 0.00 (1.3%) |
| $\omega_{\text{bats}}$ | | | | | | | 0.13 (37%) | 1.0 (62%) | 13.1 (1.3%) |
| Orthogroup B |  |  |  |  |  |  |  |  |  |
| *M2a_rel (null)* | -4071 | 20 | 2.63 | 2.59 | --- | --- |  |  |  |
| $\omega_{\text{background}}$ | | | | | | | 0.34 (63%) | 1.0 (0%) | 1.31 (37%) |
| *Clade Model C* | -4066 | 21 | 2.58 | 2.56 | 17.5 | 2.9e-5 |  |  |  |
| $\omega_{\text{background}}$ | | | | | | | 0.30 (20%) | 1.0 (54%) | 0.70 (26%) |
| $\omega_{\text{bats}}$ | | | | | | | 0.30 (20%) | 1.0 (54%) | 0.03 (26%) |
| Orthogroup C |  |  |  |  |  |  |  |  |  |
| *M2a_rel (null)* | -7145 | 36 | 2.52 | 4.78 | --- | --- |  |  |  |
| $\omega_{\text{background}}$ | | | | | | | 0.18 (24%) | 1.0 (62%) | 0.18 (14%) |
| *Clade Model C* | -7142 | 37 | 2.52 | 4.77 | 5.8 | 0.02 |  |  |  |
| $\omega_{\text{background}}$ | | | | | | | 0.23 (21%) | 1.0 (62%) | 0.23 (17%) |
| $\omega_{\text{bats}}$ | | | | | | | 0.23 (21%) | 1.0 (62%) | 0.00 (17%) |

lnL: log-likelihood; np: number of parameters; TL: tree length; K: transition/transversion rate; LR: likelihood ratio; p: p-value of likelihood ratio of alternative relative to null for each test

**Table 2.**
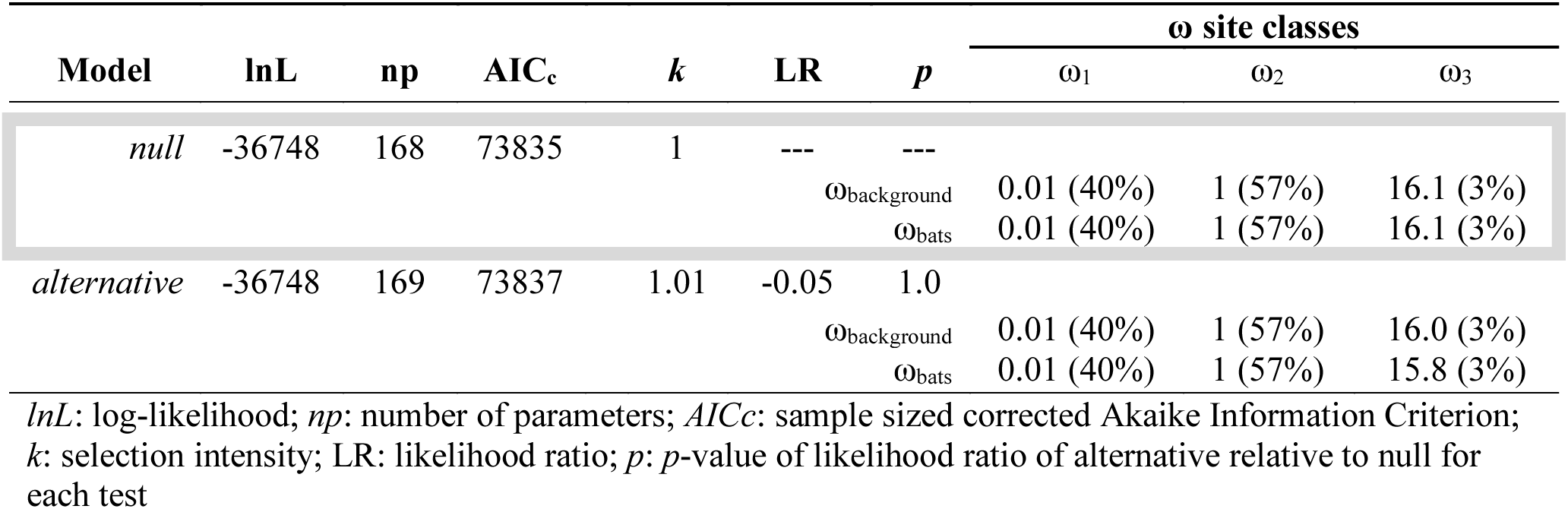
Results from RELAX analyses. Values for the site classes are ω estimates for each of the three site classes: purifying (ω_1_), neutral (ω_2_), and positive (ω_3_). The percentage values in parentheses are the proportion of sites found within that respective site class. The grey box indicates the model with the best fit, demonstrating the lowest AICc.

Chemosensory genes are among the fastest-evolving in the mammalian genome, second only to genes involved in pathogen-recognition (Yoder & Larsen 2014; Wynn et al. 2012). As the neural mechanisms of signal processing are highly conserved in vertebrates (Bear et al. 2016), the duplicative nature of these genes and the high rates of evolution likely reflect fine-tuning the detection for ever-changing environmental chemical space.

Contrary to what is seen in the gene tree as a whole, orthogroups are to be evolving differently in bats and non-bats. For some *V1r*s, bats have a higher rate and for others, horses have a higher rate (Table S3), indicating potential clade-specific adaptation of particular receptors. There were significant differences between bats and non-bats in all three orthogroups (Table 1; A: χ^2^_(1)_ = 17.5 *P* = 2.9e-5; B: χ^2^_(1)_ = 17.5 *P* = 2.9e-5; C: χ^2^_(1)_ = 5.8 *P* = 0.02). For Orthogroup A, a few sites (1.3%) were evolving at a very high rate in bats, potentially indicating adaptive selection in this group of *V1rs*. This orthogroup also showed recent duplications within *Artibeus fraterculus* leading to four detected copies. In Orthogroups B and C, bats showed low ω rates relative to the background branches for 17% and 26% of the sites, indicating strong purifying selection in bats for these genes.

Both putatively functional and pseudogenized bat *V1r*s illuminate the evolutionary processes shaping the vomeronasal system as a whole (Yohe & Dávalos 2018). The same copies of some receptors have been maintained since the ancestor of bats diverged from those of the horse or dog, as shown by both the high degree of orthology (Fig. 2), and slight differences in rates of evolution between the intact receptors of bats and those of related non-bats (Tables S3, S4). This finding bolsters the hypothesis that phyllostomid and miniopterid bats with seemingly intact vomeronasal systems retained function throughout bat diversification, while most other bat families independently lost function. Moreover, our data support the idea that all components of the vomeronasal system evolve together, resulting in an all-or-nothing pattern. Specifically, lineages with intact *V1R*s also have intact *Trpc2* (Fig. 1) and well-developed morphology, while bat families with pseudogenized *Trpc2* and/or degraded morphology tend to lack intact receptors. Together with analyses correlating high rates of *Trpc2* codon substitutions and loss of the vomeronasal brain region (Yohe & Dávalos 2018), patterns of *V1r* pseudogenization in bats highlight the consequences of relaxed selection on molecular components of the system. Finally, the phylogeny of bat *V1r* pseudogenes also reveals intact copies from the horse and dog are sometimes pseudogenized across all bats (Fig. 3), even in species with intact *Trpc2*. While these receptors are likely still relevant to the ecology of the horse and dog, their complete loss of function indicates they are no longer relevant to bats.

**Figure 3.**
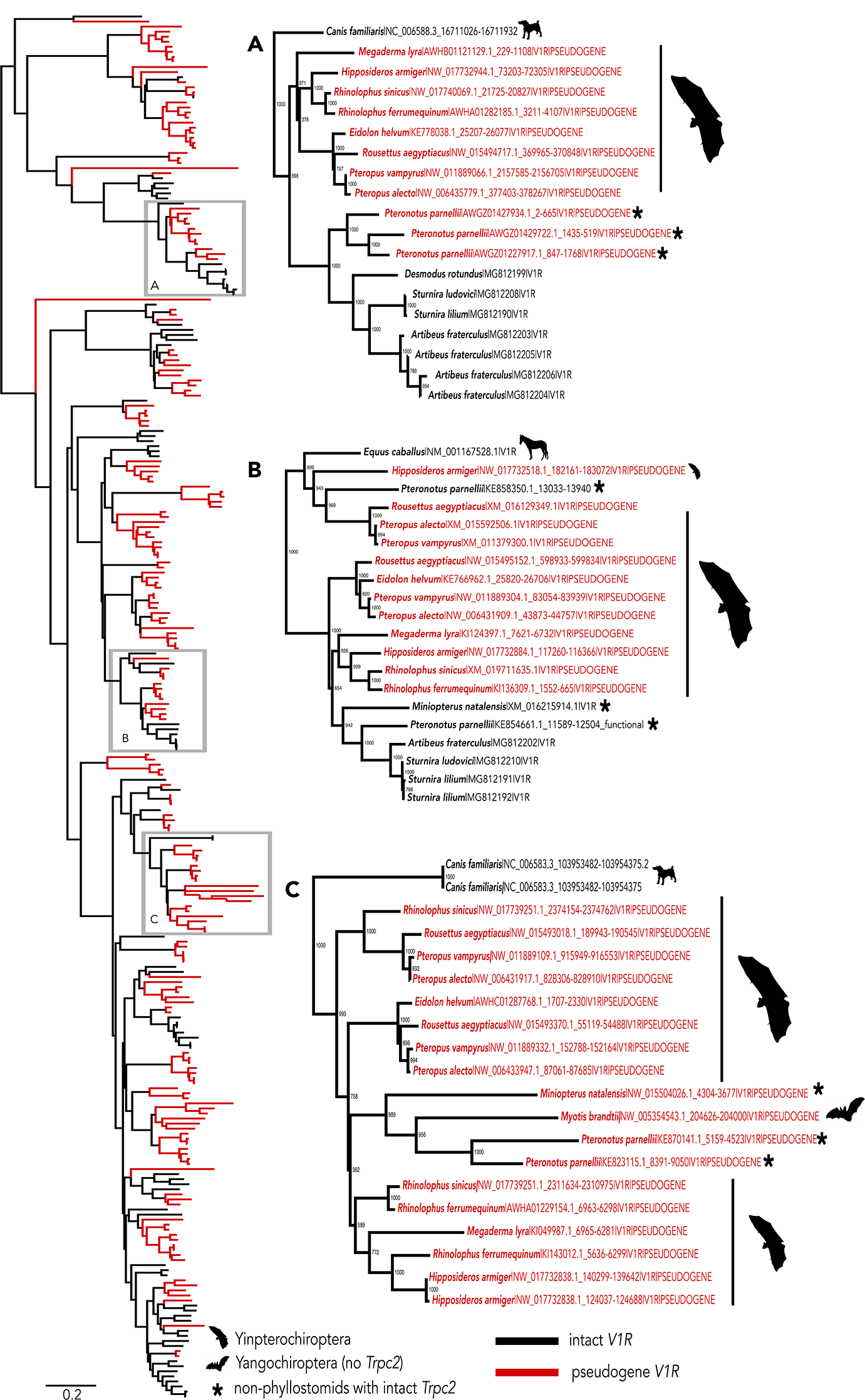
Gene tree inferred under a transitional model of nucleotide evolution of functional *V1r*s from horse, dog, and bat, as well as pseudogenes identified from all bat genomes. Horse and dog pseudogenes were not included for clarity. Red branches indicate pseudogenized genes and black indicates intact *V1r*s. Insets (A) and (B) show monophyletic groups in which the gene copy is intact in the horse or dog, and most bats with intact *Trpc2*. However, the copy has been pseudogenized in yinpterochiropteran lineages, which lack an intact *Trpc2*. Inset (C) shows a monophyletic group of genes in which the gene copy is intact in the ancestral dog, but has been lost in all bats, including species with an intact *Trpc2*. This orthogroup may be nonfunctional in phyllostomids, as there is no evidence it was expressed in the transcriptome.

Why some bats have completely lost vomeronasal function, while some have been under strong selection to retain it remains a mystery. We propose another group of receptors other than *V1r*s, such as those expressed in the main olfactory epithelium, may respond to pheromones becoming sufficient for detecting the relevant social chemical cues. While *V2r*s, the other major vomeronasal receptor gene family, were not found in the transcriptomes of these bats, the dog and cow genomes also lack *V2r*s, and these genes might not be relevant in laurasiatherians (Grus et al. 2007). In contrast to rodents, sheep and goats (also laurasiatherians) primarily use their main olfactory system for processing social chemical signals (Keller & Lévy 2012). The many genes expressed in the main olfactory epithelium, including the *major histocompatibility complex* (MHC), *trace amine-associated receptors* (*TAAR*s), and olfactory receptors, all have been shown to play a role in social chemical communication (Fortes-Marco et al. 2013; Li et al. 2013; López et al. 2014). An association between mate choice, and *MHC-*class 1 alleles and variation within *TAAR3 —*both gene families that express in the main olfactory epithelium— was recently reported for the greater sac-winged bat (*Saccopteryx bilineata*), a bat with no vomeronasal organ but with large scented glands embedded in the wing membrane (Santos et al. 2016). As odorant-binding ligands have been identified for none of the hundreds of bat olfactory receptors (Hayden et al. 2014), some may respond to pheromonal cues.

Numerous neurobiological and behavioral studies have described strong interactions between the main olfactory epithelium and vomeronasal organ in detecting and discriminating pheromonal cues and initiating the behavioral response (Fraser & Shah 2014). If the main olfactory epithelium has the potential to maintain behaviors critical to survival, then bat vomerolfaction may be redundant and susceptible to relaxed selection, explaining its frequent loss among bats.

## Material and Methods

RNA-seq libraries of the vomeronasal organ were generated for six phyllostomid species (Table S1). Reads underwent quality control (Table S2), were assembled using Trinity v. 2.2.0 (Grabherr et al. 2011), and screened for chimeric transcripts. Vomeronasal tissue was validated by identifying the tissue-specific ion channel *Trpc2 β* isoform transcripts (GenBank: MH010883-MH010888). Vomeronasal receptors were identified in the six new bat transcriptomes, and 16 published genomes (14 bats including the vampire bat, and the horse and dog) through a modified pipeline (Hayden et al. 2010) that implements a hidden Markov model algorithm to search for similar sequences using HMMER v. 3.12b trained from *V1r* sequence motif profiles (Eddy 2010). Sequences were aligned for intact receptors only, and then for intact and pseudogenized receptors. The best-fit model of evolution was estimated for both alignments using ModelOMatic v. 1.01(Whelan et al. 2015), and maximum likelihood gene trees were inferred from each alignment. Orthogroups were determined using the program UPhO (Ballesteros & Hormiga 2016). Rates of molecular evolution (ω) were estimated for bat and non-bat branch classes using Clade Model C in PAML v. 4.8 (Yang 2007) and RELAX (Wertheim et al. 2014).

## Acknowledgements

We thank Nancy Simmons at the American Museum of Natural History for help with tissue collection in Belize. We thank Centro de Ecología y Biodiversidad CEBIO, Erika Paliza, Miluska Sánchez, Jorge Carrera, Edgar Rengifo Vásquez, Harold Porocarrero Zarria, Jorge Ruíz Leveau, Jaime Pecheco Castillo, Josh Potter, Carlos Tello, Fanny Cornejo, and Fanny Fernández Melo for making tissue collection in Peru possible. We thank the Beijing Genome Institute for sequencing. NSF-DEB 1442142 to LMD and SJR, and the NSF Graduate Research Fellowship, NSF-DEB 1701414, American Society of Mammalogists Grant-in-Aid, Society for the Study of Evolution Rosemary Grant, Sigma Xi Grant-in-Aid, The Explorer’s Club, and the Tinker Foundation awards to LRY funded this study. The Indiana University Mason server funded by NSF-DBI 1458641 and the CIPRES Science Gateway provided the computational resources for the transcriptome assemblies and phylogenetic inference.

## Authors Contributions

LRY conceived the idea, collected and analyzed the data, and wrote the manuscript. LMD supported and guided material and data collection, as well as analyses, and co-wrote the manuscript. KTJD assisted in methodology and SJR provided assistance with sample collection. All authors edited the manuscript.

